# Polymer brushes immersed in solvent molecules at thermal equilibrium: A theoretical approach

**DOI:** 10.1101/404103

**Authors:** Mike Edwards

## Abstract

By means of density functional theory (DFT), influence of solvent molecules on polymer brushes is investigated. Osmotic pressure of solvent molecules gives rise to a stronger stretching of the brush chains in perpendicular direction. This suggests that the osmotic pressure of solvent molecules is a driving force in increasing thickness of brush layer.

## INTRODUCTION

Polymer brushes are formed when linear macromelecular structures are densely grafted to a surface. Steric repulsion among monomers stretches the chains in the perpendicular direction. This is a means of tuning surface properties for a wide variety of applications. Among them, the stabilization of colloidal dispersions, microfluidic devices, coatings against fouling and corrosion and lubrication are of great importance^2^. Glycol on the outside of cell membranes as well as aggregan in synovial fluids of mammalian joints, are examples of brush-like structures in biological systems^2^. Polymer brushes have been subject of theoretical, numerical and experimental investigations which interested readers may read a recent review and the references therein^4^.

In most practical situations, brush chains are immersed in solvent molecules. Despite many theoretical studies on dry brushes, there is a lack of investigations on brushes in solvent. To this end, here I tackle this problem by means of density functional theory (DFT). I do not take the excluded volume interactions between solvent molecules and monomers as well as between solvent molecules. Therefore, the results are valid in solvents with low density. However, as the excluded volume interactions are screened in higher concentrations, the results can be extended well to larger solvent concentrations.

### DENSITY FUNCTIONAL THEORY (DFT) METHOD

#### Ideal gas as solvent molecules

In statistical mechanics, a system of *N* non-interacting particles with mass *m* are of great importance^3^. These particles have only linear momentum and the Hamiltonian of this system is given as follows,

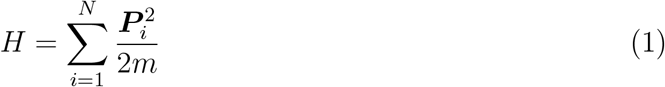

where ***P***_*i*_ is linear momentum vector of i-th particle. Since, brushes are usually in room temperature, we can consider classical particles with a good approximation. The classical partition function is given as follows^3^,

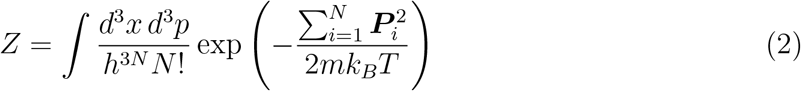

Where *h* = 6.63 × 10^−34^*J*.*s* is the Planck’s constant and *k*_*B*_ = 1.38 × 10^−23^*J K*^−1^ the Boltzmann’s constant. The linear momentum of different particles in different directions are uncorrelated. This means that the partition function can be obtained by using Gaussian integral solution as follows,

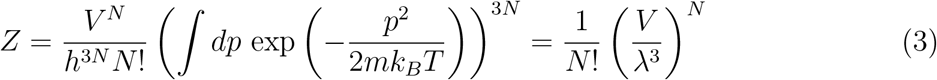

Where *λ* = *h/*(2*πmk*_*B*_*T*) ^1*/*2^ is the de-Broglie thermal wavelength and it is a measure of particle size in a certain temperature. For instance, electrons with rest mass 9.1 × 10^−31^*kg* at room temperature (300*K*) have a thermal wavelength *λ*_*e*_ ≈ 4.3*nm* while protons and neutrons with rest mass 1.67 × 10^−27^*kg* have thermal wavelength *λ*_*p*_ = *λ*_*n*_ = 0.1*nm*^3^. Therefore, it appears that heavier particles occupy smaller portion of space. However, it has been confirmed that quantum corrections for electrons are necessary as they become important.

The Helmholtz free energy of an ideal gas is calculated as follows,

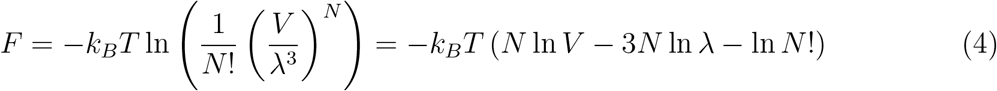

Nevertheless, the last term does not seem to be an straight-forward term in powers of *N*. That’s why the Stirling formula is used as a good approximation to *N* !. Stirling argued that 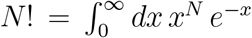. This integral is not exactly solvable with known methods. Stirling proposed that the integrand become fitted to a Gaussian distribution function *A* exp −(*x* − *N*) ^2^*/a*^2^ up to the second order. This way the Stirling’s approximation to *N* ! has been obtained which is given as follows,

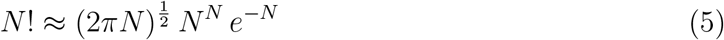

By inserting the Stirling formula into Eq. (4) and simplifying it, we get the following expression for the Helmholtz free energy per unit volume,

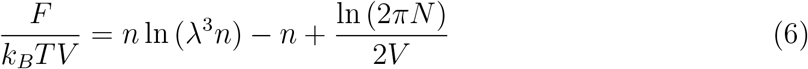

Where *n* = *N/V* is particle density. The third term can be ignored as it is small (ln *N/V*) compared to terms. The osmotic pressure per unit volume is given as follows,

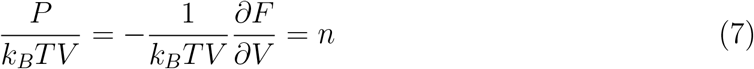

This is the *thermal equation of state* of an ideal gas. The entropy per unit volume is given as follows,

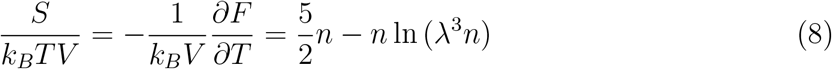

Having the equation of state and the entropy we can calculate the internal energy per unit volume of an ideal gas that is given as follows,

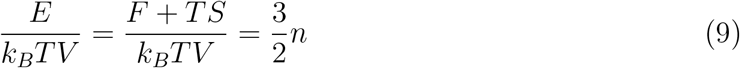

Now let me assume an ideal gas at thermal equilibrium. Since, number of particles is fixed, I need to transform to the grand free energy as follows,

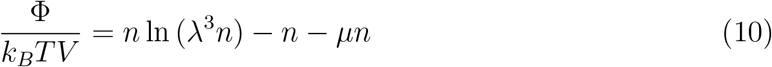

where the last term is a constraint to fix number of particles and *µ* is the corresponding Laggrange multiplier. However, *µ* is called *chemical potential* as it is a measure of changing the free energy under variation of particle number. The equilibrium density is obtained through solving the following Euler-Laggrange equation,

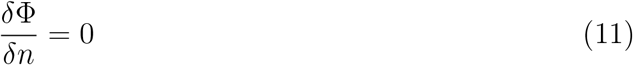

This equation gives us the equilibrium particle density as,

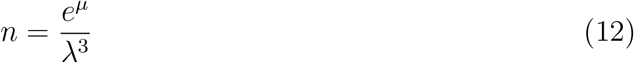

This result tells us that if *µ* = 0, the particle density at thermal equilibrium becomes *λ*^−3^. This confirms the idea that each particle occupies a portion of space equal to *λ*^3^.

### Polymer brushes

The grand free energy of a polymer brush is made by balancing the leading order term of the entropic elasticity of polymer chains in the fixed length ensemble as well as the excluded volume interactions among monomers^6–8^. The fixed number of monomers is applied as a constraint via a third term with chemical potential as the Laggrange multiplier. The polymer chains are grafted to a substrate located at *z* = 0 and the chains stretch in perpendicular direction that is z-direction. The grand potential functional of polymer brushes is given as follows,

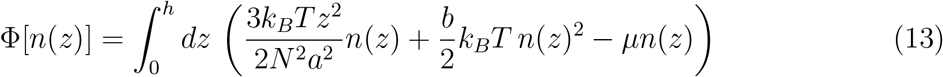

where *h* is the brush height, *N* number of monomers per chain, *a* the Kuhn or segment length which gives us size of a monomer, *b* = (1/2) ∫ d^3^*r*(1 − exp (−*U*(*r*)/*k*_*B*_*T*)) the second Virial coefficient which defines the binary correlations among monomers and *µ* the chemical potential. The second Virial coefficient for hard spheres of diameter *a* is equal to (2*π/*3)*a*^3^ and for the Lennard-Jones particles with binary potential *U* (*r*) = 4*∈*[(*a/r*)^1^2 − (*a/r*)^6^] equal to the following series,

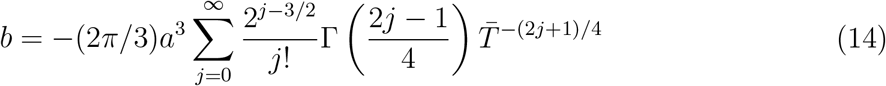

Where 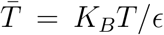 is dimensionless temperature and Γ is the gamma function. For 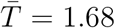, the second Virial coefficient of Lennard-Jones particles converges to *b* = 4.19767. There are three unknown variables in Eq. (13), the brush height *h*, the monomer density profile *n*(*z*) and the chemical potential *µ*. To find them we need three equations which are given as follows,

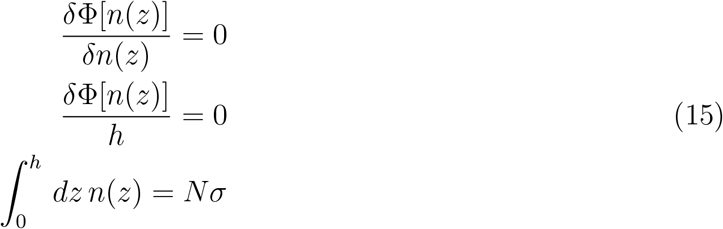

with *σ* number of grafted chains per surface area i.e. grafting density. By solving the above set of coupled equations we get the following results for unknown variables,

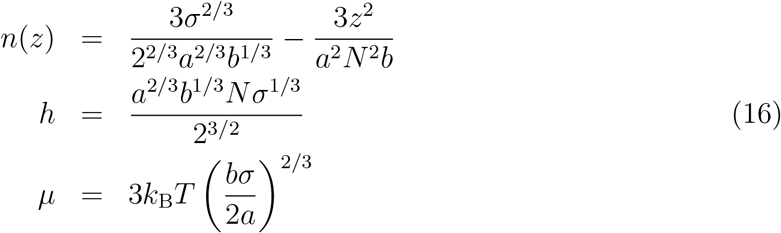

By inserting the above results into Eq. (13), one finds the following grand potential energy in terms of molecular parameters of brush,

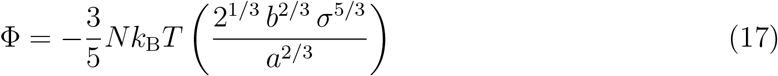

## RESULTS

### Polymer brushes immersed in solvent molecules

Let us assume a polymer brush covered surface at *z* = 0 which is immersed in a solvent constrained between the brush-covered surface and a flat surface at *z* = *D*. To avoid the surface effects, I assume *D* ≫ *h*. Here, I am interested in influence of solvent molecules on the equilibrium properties of brush. To this end, I divide the channel into two regions; (i) inside brush and (ii) outside brush. The grand potential energy for both regions are given as follows,

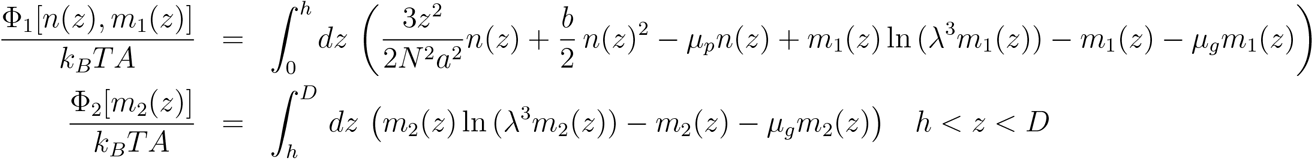

where I have used the grand free energy of brush Eq. (13) and grand free energy of solvent Eq. (10). It is worth noting here that, the chemical potential is the same throughout the channel as number of solvent molecules is fixed in it. However, solvent density profiles are assumed to be different in each region. This is a key point to take the solvent particle exchange between two regions into account. There are six unknown variables in this problem; the brush height *h*, brush density profile *n*(*z*), solvent density profile inside *m*_1_(*z*) and outside brush *m*_2_(*z*), brush chemical potential *µ*_*p*_ and solvent chemical potential *µ*_*g*_. To find the values of these variables at thermal equilibrium, one has to solve the following six coupled equations simultaneously,

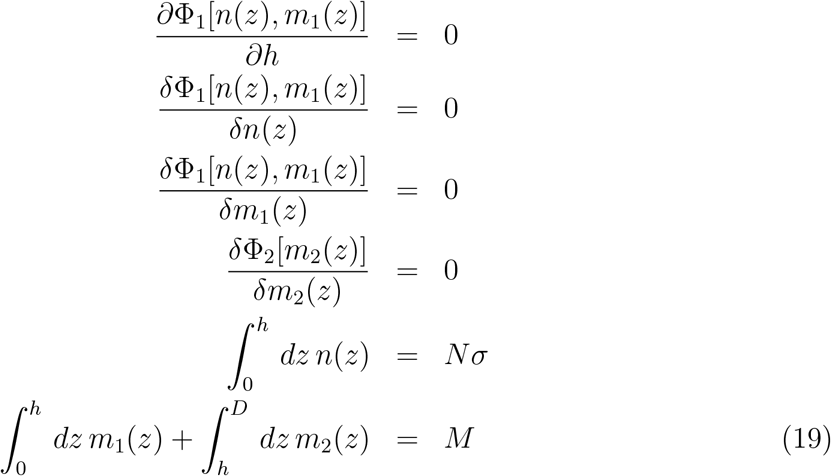

where *M* is total number of solvent molecules per unit area. By solving the above set of coupled equations one obtains the following results,

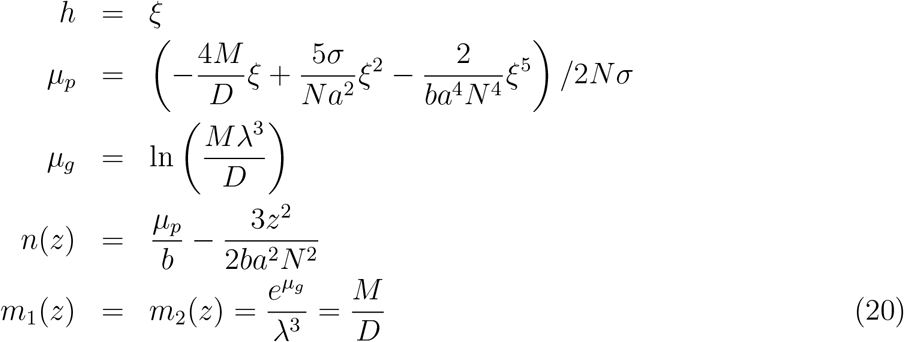

where *ξ* is a the solutions of the following sixth degree equation,

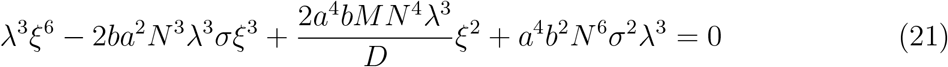

One can not analytically solve Eq. (21) with known methods. That is why, hereafter, I utilize a numerical package in the Wolfram Mathematica i.e. *RootFinding* to find approximate solutions of Eq. (21). Let me choose *N* = 30, *a* = 1, *b* = 4.19767, *σ* = 0.1 as molecular parameters of brush and *λ* = 1 and *M* = 0.1 as the one for solvent and set *D* = 50. Basically, here I choose *a* = *λ* = 1 to make size of monomers and solvent molecules equal. Furthermore, I choose *σ* = *M* = 0.1 to initially have a brush regime as well as a low density of solvent molecules.

As one can see in Fig. (1), the osmotic pressure of solvent molecules strongly stretches brush chains in perpendicular direction. Up to this approximation which the excluded volume interactions between monomer-solvent as well as solvent-solvent are ignored, while, the excluded volume interactions between monomer-monomer has been taken into account, stretch of brush chains is captured, however, solvent density does not change in both regions.

**Figure 1:**
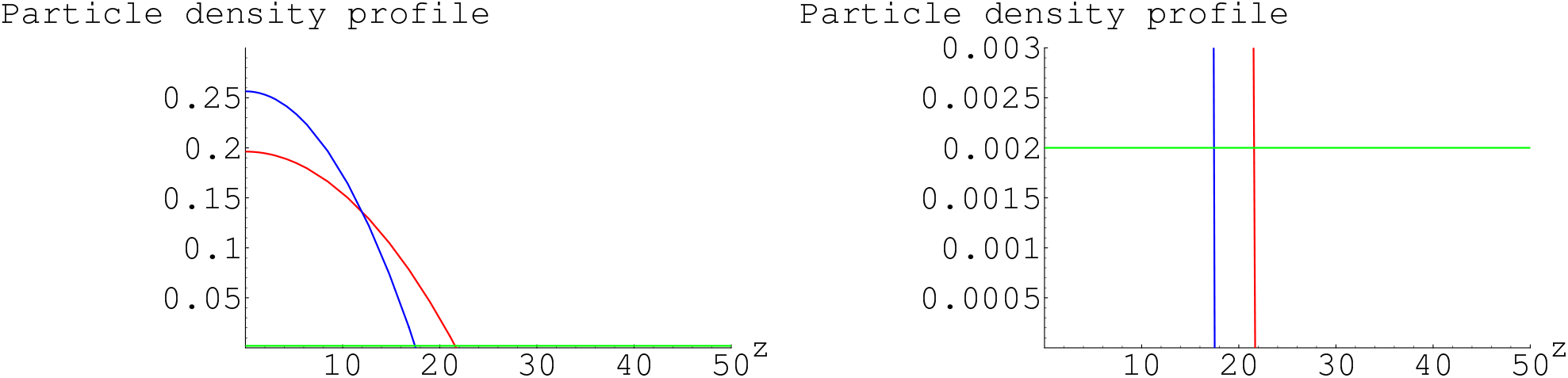
Left: Density profile of brush immersed in solvent molecules (Red), density profile of solvent (Green) and density profile of dry brush (Blue). Right: A zoomed view on density profile of solvent molecules.

## CONCLUSIONS

The osmotic pressure of solvent in polymer brushes is very challenging problem in theoretical polymer, soft matter and biological science. There has been a long-standing discussion among theoreticians about whether the osmotic pressure of solvent molecules inside a brush, leads to chains further stretching. In particular, about the Polyelectrolyte brushes this had remained unknown whether swelling of Polyelectrolyte brushes is due to osmotic pressure of the counter-ions inside brush or, otherwise, due to electrostatic repulsion between monomers. Here, I demonstrated theoretically that the osmotic pressure of solvent molecules have a considerable contribution (As chains stretch to around 25% of their initial length) on chains stretch. Furthermore, I chose molecular parameters such that absence of the solvent-monomer and solvent-solvent excluded volume interactions does not influence the results. Therefore, one can say that at low density of solvent molecules, the osmotic pressure of solvent is dominant factor causing chain stretch. Nonetheless, the excluded volume interactions become screened upon increasing concentration. This means that even at larger concentrations, the osmotic pressure is considerably important. This research can be extended to systems with excluded volume interactions as well as systems with long-ranged electrostatic interactions.

## Notes

### Competing Interest Statement

The authors have declared no competing interest.

### Summary of Updates

The corresponding author name and his E-Mail address is changed. The template of the manuscript is changed. In the conclusion section, the citation to an unpublished work is erased.

